## Supplement for "SUNi mutagenesis: scalable and uniform nicking for efficient generation of variant libraries"

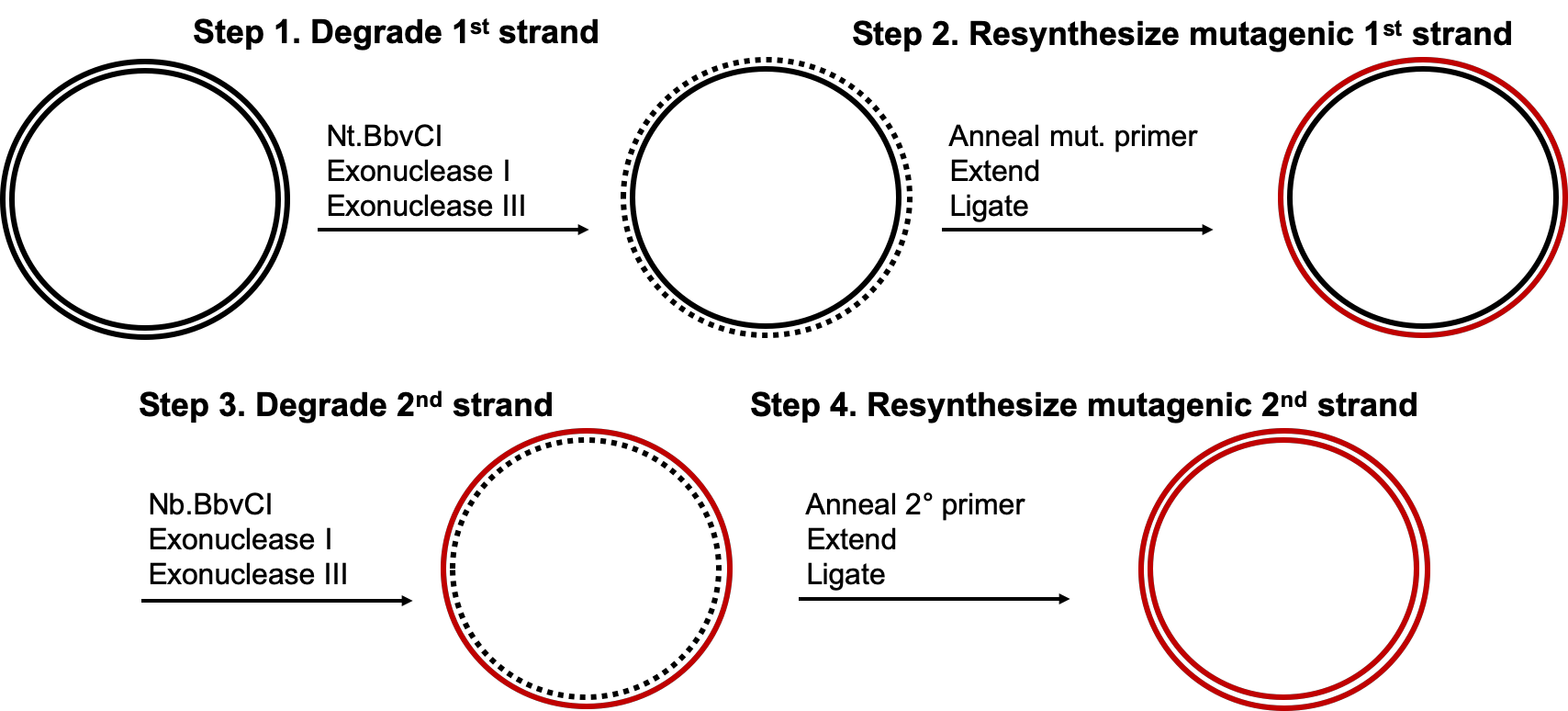

**Supplementary Figure 1. Schematic overview of the four main steps of nicking mutagenesis**

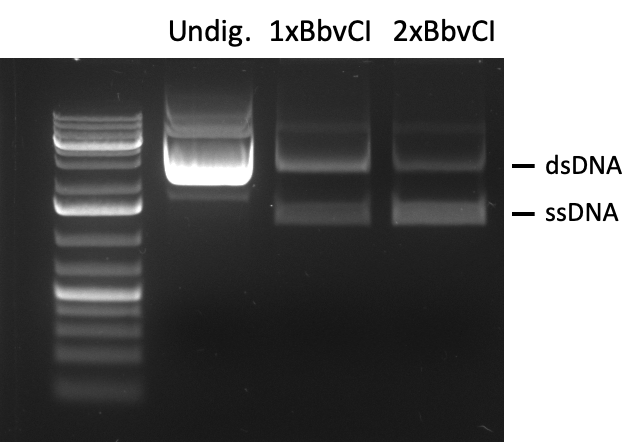

**Supplementary Figure 2. Two BbvCI sites improves digestion efficiency**

Plasmids with either one or two BbvCI sites were digested with Nt.BbvCI, exonuclease I and exonuclease III, per the original nicking protocol, and digestion products were visualized on an agarose gel.
