## Supplementary protocol for "SUNi mutagenesis: scalable and uniform nicking for efficient generation of variant libraries"

**SUNi mutagenesis wet lab protocol.**

**Reagents**

- Mutagenic SUNi primers and secondary primer (that binds outside the mutated region and to the opposite strand as the mutagenic primers).
- Enzymes (all purchased from NEB; diluent for all enzymes is 1X NEB CutSmart Buffer):
  - T4 Polynucleotide Kinase (10 U/µL) [2µL]
  - Nt.BbvCI (10 U/µL) [1µL]
  - Nb.BbvCI (10 U/µL) [1µL]
  - Exonuclease III (100 U/µL) [2µL]
  - Exonuclease I (20 U/µL) [2µL]
  - Phusion High-Fidelity DNA Polymerase (2 U/µL) [2µL]
  - Taq DNA Ligase (40 U/µL) [10µL]
  - DpnI (20 U/µL) [2µL]
- 50 mM DTT
- Electroporation cuvettes
- Electrocompetent E. Coli

**Protocol**

1. Phosphorylate mutagenic SUNi and secondary primers.

|  | Mutagenic oligo mix | Secondary primer |
| --- | --- | --- |
| oligos | 20µL @ 10µM | 7µL @ 100µM |
| T4 PNK Buffer | 2.4µL | 3µL |
| 10mM ATP | 1µL | 1µL |
| T4 PNK (10 U/µL) | 1µL | 1µL |
| H2O | -- | 18µL |

- Incubate 37° for 1 hour

2. Prepare ssDNA template

|  | Per sample |
| --- | --- |
| Plasmid dsDNA (0.76 pmol) | X |
| 10x CutSmart Buffer | 2µL |
| Exonuclease I (20 U/µL) | 1µL |
| 1:10 dil. Exo III (10 U/µL) | 1µL |
| Nt.BbvCI  (10 U/µL) | 1µL |
| H2O to 20µL final vol. | 15µL - X |

- Incubate at 37° for 1 hour; 80° for 20 minutes; 10° hold.

3. Mutagenesis

|  | Per mutagenic sample |
| --- | --- |
| H2O | 26.7µL |
| 5x Phusion HF Buffer | 20µL |
| 1:500 diluted Mutagenic oligo (step 1) | 4.3µL |
| 50mM DTT | 20µL |
| 50mM NAD+ | 1µL |
| 10mM dNTPs | 2µL |
| Phusion HF Polymerase (2 U/µL) | 1µL |
| Taq DNA LIgase (40 U/µL) | 5µL |

- PCR program: 98° for 2 minutes; 5 cycles of [98° for 30 s, 55° for 45 s, 72° for 7 min], 10° hold.

4. Column purification
   - Purify the previous reaction with MinElute (New England Biolabs) columns and elute in 15 µL of NFH_2_O into DNA LoBind tubes (Eppendorf)
5. Degrade template strand

|  | Per sample |
| --- | --- |
| Eluate from step 4 | 14 µL |
| 10x CutSmart buffer | 2 µL |
| Exonuclease I (20 U/µL) | 2 µL |
| 1:50 diluted Exo III (2 U/µL) | 1 µL |
| 1:10 diluted Nb.BbvCI (1 U/µL) | 1 µL |

- Incubate 37° for 1 hour; 80° for 20 minutes, 10° hold.

6. Synthesize 2^nd^ mutagenic strand

|  | Per tube |
| --- | --- |
| Reaction from step 5 | 20 µL |
| H_2_O | 27.7µL |
| 5x Phusion HF buffer | 20µL |
| 1:20 diluted phosphorylated secondary primer | 3.3µL |
| 50mM DTT | 20µL |
| 50mM NAD+ | 1µL |
| 10mM dNTPs | 2µL |
| Phusion HF Polymerase (2 U/µL) | 1µL |
| Taq DNA Ligase (40 U/µL) | 5µL |

- PCR program: 98° for 30 seconds, 55° for 45 seconds, 72° for 10 minutes, 45° for 20 minutes; 10° hold.
- [Safe stopping point; store reactions in 4° overnight]

7. DNA cleanup
   - Add 2 µL of DpnI to each reaction and incubate 37° for 1 hour.
8. Column purification
   - Purify the previous reaction with MinElute (New England Biolabs) columns and elute in 6 µL of NFH_2_O into DNA LoBind tubes (Eppendorf)
9. Bacterial electroporation
   - Use 1.5 uL of purified plasmid to transform electrocompetent cells (for example 10-beta Electrocompetent E. Coli (New England Biolabs))
